# An efficient stochastic steering strategy of magnetic particles in vascular networks

**DOI:** 10.1101/2023.02.23.529635

**Authors:** Kejie Chen, Rongxin Zhou, Xiaorui Dong

## Abstract

One of the primary challenges of magnetic drug targeting is to achieve efficient and accurate delivery of drug particles to the desired sites in complex physiological conditions. Though a majority of drugs are delivered through intravenous administration, until now, the kinematics and dynamics of drug particles influenced by the magnetic field, vascular topology and blood flows are still less understood. In this work, a multi-physics dynamical model which captures transient particle motions in both artificial and *in vivo*-like 3D vascular networks manipulated by the time-varying magnetic field is developed. Based on the model, it is found that particles which perform a random walk with correlated speed and persistence (RWSP motion) inspired by the migratory motion of immune and metastasis cells have higher mobility and navigation ability in both 2D and 3D tree-like and web-like networks. Moreover, to steer particles to perform the efficient RWSP motion, a stochastic magnetic steering strategy which uses time-varying gradient magnetic field is proposed. Parameters of the steering strategy is optimized and the capability of controlling particles to achieve fast spreading and transport in the vascular networks is demonstrated. In addition, the influence of heterogeneous flows in the vascular networks on the particle steering efficiency is discussed. Overall, the numerical model and the magnetic steering strategy can be widely used in the drug delivery systems for precise medicine research.

## 1. Introduction

Magnetic drug targeting (MDT) is a novel, minimally invasive and efficient method to actively deliver drugs to lesion sites, in order to improve drug therapeutic efficacy and mitigate potential adverse effects [1–3]. In MDT, synthesized magnetizable particles which carry drug molecules are injected into the blood streams or airways, and then steered to the desired location by external magnetic fields. The MDT concept was first demonstrated by Zimmermann et al. in the 1980s, who used magnetic erythrocytes for delivering cytotoxic drugs [4]. Since then, many research groups and start-ups around the world have been devoting lots of efforts in developing magnetic drug carriers, magnet systems and control strategies for successful and efficient drug targeting.

Nowadays, the production of magnetic drug carriers with desirable composition [5], size [6] and shape [7] for MDT have been well-established. Metal [8], polymer [9], gel [10] and hybrid [11] particles of various shapes such as spherical, quasi-cubical and rod can be fabricated [12–14] to carry different types of drug molecules [15], peptides [16] and aptamers [17]. However, a full clinical *in-vivo* application of MDT is still subject to numerous open challenges. One of the most critical challenges is how to efficiently concentrate drug particles to the target sites, especially to the deep tissues within the body.

Drug particles are usually injected into the body through intravenous [18], oral [19] or nasal [20] administration. Many factors, such as the biological fluid flow [21], vascular or airway structures [22], magnetic and surface characteristics of particles [23], magnetic field strength and gradients [24], can influence particle motion and retention. Previous works decipher the drug delivery mechanisms and address the drug targeting challenge mainly from two ways. 1) For an open-loop control system of magnets, numerical models are performed to optimize the properties of particles and magnets; The design of drug delivery systems are verified by both *in-vitro* and animal experiments [25–29]; 2) Advanced control and navigation strategies guided by the real-time imaging of particle locations and environmental conditions are included to improve the targeting accuracy [30–33].

The open-loop control of magnetic particles is relatively simple and robust, which has been the most widely used strategy so far. By numerically computing the forces on the magnetic particles and simulating particle dispersion and penetration through tissue layers, the effects of particle properties, magnetic field, tissue properties and fluid flows on the drug targeting efficacy can be understood [25–29, 34–36]. For example, it has been shown that increasing particle diameter, magnetic field strength improves delivery efficacy in various flow conditions including steady state flow, non-Newtonain blood flow, and pulsatile arterial flow [26–29]. Ring-shape magnet, cylindrical magnet, Halbach array are found to be efficient in capturing drug particles [24]. Moreover, the rotating or pulse magnetic fields can reduce the formation of particle aggregates and enhance the passage of particles through the anatomical barriers like blood-brain barrier [37]. These mechanistic studies and conclusions have provided valuable insights in designing novel drug carries and magnet systems, and have greatly improved the MDT efficacy in the experiments. However, most current studies only consider particles moving in the cylindrical or Y-shape bifurcated vessels, but neglect the realistic and complex anatomical geometries and the physiological boundary conditions. The dynamics of drug particles in the *in-vivo* relevant conditions is still rarely known.

To better navigate in the vascular networks (or gastrointestinal and respiratory tracts), advanced microrobotic drug carriers assisted by the imaging feedback control systems are developed. Drug particles are visualized using imaging techniques [38] such as magnetic resonance imaging (MRI) [39], ultrasound imaging [40] and computed tomography (CT) [41]. Automated control schemes of magnets are developed to use real-time imaging information to guide the motion of magnetic particles. Nevertheless, the imaging feedback control is limited by the imaging resolution and the low frame rates. Particles less than 100 *µm* can hardly be recognized by most imaging techniques [33], though they are widely used in MDT because the small size facilitate circulating without causing blockage. In addition, the imaging frame rate is usually lower than 50 *Hz*, so drug particles can easily swim out of the view field and get lost between two consecutive frames, especially in the fast and heterogeneous flow conditions [42]. Therefore, it is of great necessity to develop new steering strategy of these tiny drug particles and improve the drug delivery and targeting ability in tortuous vascular networks.

In this work, we developed a multi-physics numerical model to study the dynamics of magnetic particles in 2D and 3D vascular networks using an Euler-Lagrange approach, where the network boundaries, magnetic field and base fluid are treated as the continuum phase and particles are considered as the discrete phase. An open-loop stochastic steering strategy of magnetic particles in the vascular networks inspired by the efficient migratory motion of immune cells or metastasis cells is proposed. The performance and efficacy of the stochastic steering strategy in the 2D, 3D artificial branching networks and 3D *in-vivo* like networks are demonstrated using the multi-physics model. Specifically, a few recent studies have found that immune cells including T cells and dendritic cells and cancer cells adopt an efficient search and navigation strategy through the persistence-speed coupling mechanism [43–45]. That is, the straightness of cell movement (i.e., the persistence of a cell moving along the same direction) is an exponential function of moving speed. Our previous work have also demonstrated that a particle performing random walks with exponentially correlated speed and persistence (RWSP) achieves fast spreading and efficient searching ability in both homogeneous and porous media [46]. Therefore, in this work, we first analyzed the mobility of a particle performing cell migratory-like RWSP motion in 2D and 3D branching networks, and compared its mean square displacements (MSDs) with persistent random walk (PRW) [47] and Lévy walk [48]. Next, we designed a stochastic strategy that can steer the magnetic particles to perform the RWSP type motion. The motion of the particle controlled by the stochastic magnetic field in a 3D vascular network constructed from the *in-vivo* image is studied. Furthermore, the influence of heterogeneous flow field in the vascular network on particle manipulation and transport is discussed. Overall, the open-loop stochastic magnetic steering strategy proposed in this work can be easily implemented and used in the experiments without using expensive and cumbersome imaging equipment. Thus, it has the high competence and potential to be widely used in the noninvasive drug delivery and microrobotic surgery systems.

## 2. Materials and methods

In this section, the multi-physics model developed to study the dynamics of magnetic particles in the vascular networks (Figure 1 (a)) is introduced. Specifically, in section 2.1, particle dynamics in the 2D and 3D vascular networks, including PRW, Lévy walk and RWSP motions, are introduced. In section 2.2, the multi-physics dynamical model for simulating particle motion controlled by the magnetic field and influenced by the base flows is discussed. The reconstruction of 3D *in-vivo* like vascular networks is presented in section 2.3. And in section 2.4, the computational procedure and the pseudo-code of the multi-physics dynamical model is presented.

**Figure 1.**
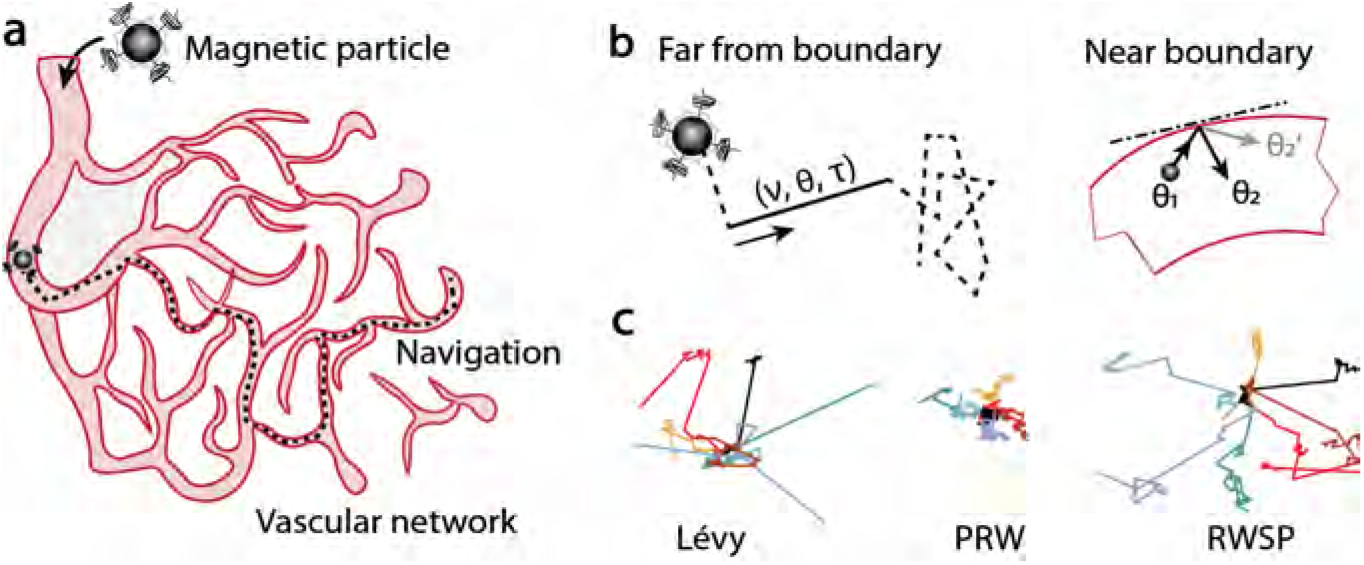
Schematics of magnetic particle dynamics in the vascular network. **(a)** Magnetic drug targeting (MDT) is achieved through magnetic particles carrying drug molecules and concentrating in the lesion site by magnetic field. **(b)** The dynamics of magnetic particles is influenced by the moving strategy and the reflection at vascular boundaries. **(c)** Example trajectories of three types of particle motions, PRW, Lévy walk and RWSP. Different colors represent different trajectories starting from the same location.

### 2.1 Particle motion in vascular networks

Moving in a complex and unknown network, a blind particle needs to adjust its moving direction and speed randomly and frequently in order to avoid being trapped in the dead ends. Previous studies have shown that particles performing random walks with correlated speed and persistence (RWSP motion) have fast spreading and searching ability in homogeneous and porous environments [43–46]. It is reasonable to hypothesize that the RWSP motion also achieves large particle mobility in vascular networks. But whether and how the topology of vascular networks influence the particle dynamics remain unknown. Thus, we first developed an agent-based Monte Carlo model to investigate the efficacy of three types of movements (i.e., PRW, Lévy walk and RWSP) in 2D and 3D vascular networks. Here, a brief introduction about the theory of PRW, Lévy walk and RWSP is given.

In general, the random motion of a particle can be simplified as consecutive straight lines. For example, until time *T*, the particle performs *n* straight movements. During a straight movement *i* (*i* = 0, 1, …, *n*), the moving speed *v*_*i*_ and the direction ***e***_*i*_ are the same. The time duration of a straight movement *τ*_*i*_ is named as the persistent time (or persistence). The particle position ***x*** at time *t, t ∈* [0, *T*] is mathematically described as

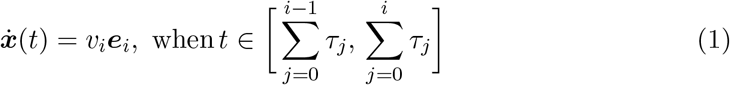

***x*** and ***e***_*i*_ are 2D or 3D vectors. For example, in the 3D space, ***x*** = [*x, y, z*] and ***e***_*i*_ = [*e*_*ix*_, *e*_*iy*_, *e*_*iz*_]. *e*_*ix*_, *e*_*iy*_ and *e*_*iz*_ are random numbers uniformly distributed in [*−*1, 1]. Then |***e***_*i*_| is normalized to one. *v*_*i*_ and *τ*_*i*_ are random numbers which determines the type of particle movements.

#### (1) Persistent random walk (PRW)

PRW and its governing differential equation (telegrapher’s equation) was proposed to resolve the paradox of the infinite speed in the heat and diffusion equations almost two century ago. In recent decades, it has been found that many nature processes such as cell migration and pollutant transport also follow a PRW process. The fundamental characteristics of a PRW process is that *v*_*i*_ and *τ*_*i*_ follow two independent exponential distributions. Specifically, 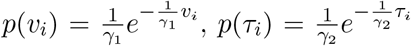where *γ*_1_ (mean velocity) and *γ*_2_ (mean persistent time) are two constant parameters [45]. When particles perform PRW type of motion, they are ballistic in the short time scale (*t < γ*_2_), and becomes diffusive at the long time scale (*t > γ*_2_) with a diffusion coefficient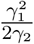

#### (2) Lévy walk

The theory of Lévy walk was originally developed to study the anomalous diffusion in turbulent flows where particles spread faster than normal diffusion. Based on the Lévy stable laws and the physical requirement of particles having finite speed, Lévy walk model considers the power law tails and diverging second moments of the straight movements of the particles. Specifically, *v*_*i*_ still follows an exponential distribution as 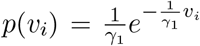But *τ*_*i*_ follows a (truncated) long tail distribution, which is represented as 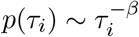, where the parameter *β* satisfies *β ∈* (1, 3) [46]. When *β* is close to 1, the particle motion is close to ballistic. When *β* is close to 3, the particle motion is close to normal diffusion. It has been shown that many self-propelled motions, such as bacteria swimming and human travels, which can take the energy from environments and convert to their own motions follow a Lévy walk type of motion.

#### (3) Random walk with correlated speed and persistence (RWSP)

Several recent studies have found that highly migratory cells such as immune cells and metastasis cells do not follow a PRW or a Lévy walk type of motion. Their moving speed *v*_*i*_ and directional persistence *τ*_*i*_ are positively correlated, such that fast cells are more likely to maintain straight movement and reach to a distant position. Specifically, *v*_*i*_ follows an exponential distribution as 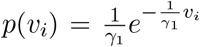 satisfies 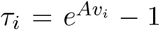 where *A* is a constant parameter [46]. Several studies including ours have shown that the RWSP type of motion greatly increase particle mobility, target searching ability and efficiency [43–46].

Simulated trajectories of the three types of motions in the homogeneous space is shown in Figure 1 (c). The mobility of the particles performing RWSP motion is the largest. Many long straight movements greatly increase the MSD and searching efficiency of the particles. The particle mobility of Lévy walk is larger than PRW. Note that, there are other types of random motions such as Brownian diffusion, correlated random walk, run and tumble dynamics. These motions are not discussed in this study because they have been proven to be less efficient than PRW or Lévy walk [49].

The particle mobility is qualified by the ensemble-averaged MSD, which is mathematically expressed as

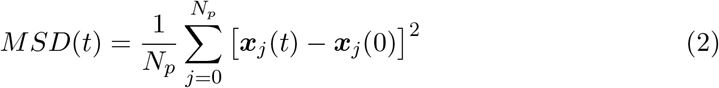

Where *N*_*p*_ is the total number of particles. *N*_*p*_ = 10^4^ in this study.

In the vascular networks, the straight movements of particles are interrupted by vessel boundaries (Figure 1 (b)). In this study, a slippery reflective boundary condition is applied. Specifically, when the particle encounters the boundary, it will automatically select a new random moving direction pointing toward the inner side of the vessel.

The probability of selecting *θ*_2_ or 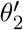 is assumed to be the same. The penetration of particles through the vascular walls is not considered in this study, though the small-size particles (nanoparticles) can potentially be engulfed by vascular endothelial cells [50], or penetrate through the vascular walls and diffuse into the surrounding tissues in the *in vivo* conditions [27].

### 2.2 Multi-physics dynamical model

Magnetic particles transport in the vascular networks is governed by several forces, including the magnetic force, hydrodynamic drag force, buoyancy, gravity and thermal kinetics. The magnetic force and hydrodynamic drag force are two dominant forces.

For particles with a diameter larger than 40 *nm*, the thermal kinetics (Brownian motion) is less influential based on the criterion developed by Gerber et al. [51], so it is neglected. Other factors are approximately an order of magnitude smaller than the dominant forces, and they are also ignored. Thus, the governing equation of particle motion is expressed using Newton’s law as [52]

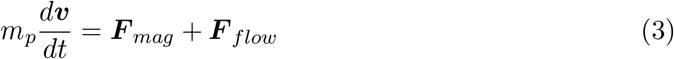

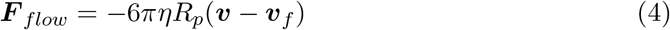

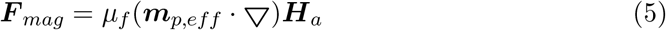

Where the inertial term of micro/nano particles 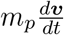 is small and can be neglected. ***F*** _*mag*_ is the magnetic force, and ***F*** _*flow*_ is the hydrodynamic drag force. *η* is the dynamic viscosity of the surrounding fluid. In this study, the surrounding fluid is assumed to be water. *η* = 10^*−*3^*kg/m*^*−*1^ *s*^*−*1^ is used. *R*_*p*_ is the hydrodynamic radius of the particle, υ is the particle velocity, υ_*f*_ is the velocity of the fluid, *µ*_*f*_ is the magnetic permeability of the transport fluid (i.e., *µ*_*f*_ = *µ*_0_ = 4*π ×* 10^*−*7^ *H/m*), ***m***_*p,eff*_ is the effective dipole moment of the particle, and ***H***_*a*_ is the intensity of the magnetic field at the center of the particle. In this study, single core, spherical Fe_3_O_4_ magnetic particles whose radius *R*_*p*_ is 5 *µm* are studied [25–27].

To determine the magnetic force, a linear magnetization model with saturation is used [52]. Specifically, the effective dipole moment of the particle is calculated as ***m***_*p,eff*_ = *V*_*p*_***M*** _*p*_, where *V*_*p*_ is the particle volume, and ***M*** _*p*_ is the magnetization of the particle. Below saturation, ***M*** _*p*_ satisfies

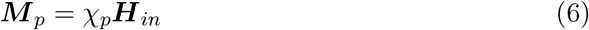

Where *χ*_*p*_ = *µ*_*p*_*/µ*_0_ *−* 1 is the susceptibility of the particle, *µ*_*p*_ is the magnetic permeability of the particle (i.e., *µ*_*p*_ = 4.1*µ*_0_) [25–27], and ***H***_*in*_ is the magnetic field intensity inside the particle. ***H***_*in*_ is acquired by solving the magnetostatic boundary value problem for a spherical particle in the fluid [52]. Specifically, the fields inside and outside the particle represented by scalar potentials are expressed as

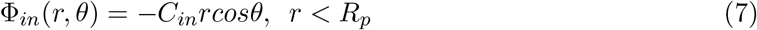

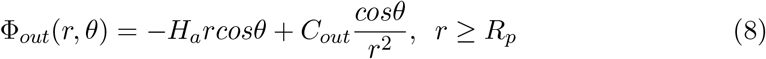

Where *C*_*in*_ and *C*_*out*_ are the magnitudes of the magnetic field intensity inside and outside the particle. *C*_*in*_ and *C*_*out*_ are obtained based on the solving the following particle boundary problem. At the particle boundary *r* = *R*_*p*_, the potentials and the normal components calculated from the fields inside and outside the particle are the same. Mathematically, it satisfies

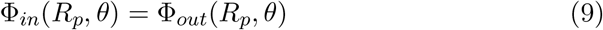

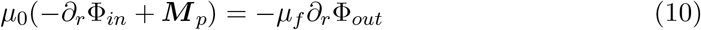

By combining Eqs. (7)-(10), the magnitude of the “equivalent” point dipole moment is expressed as [52]

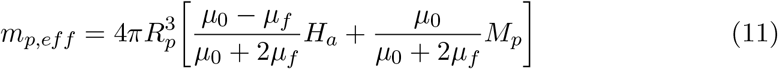

Substituting *M*_*p*_ = *χ*_*p*_*H*_*in*_ = *C*_*in*_ to get

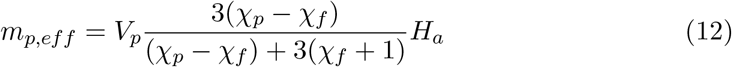

Thus, the magnetic force acting on the particle is expressed as

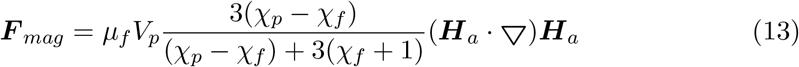

The fluid velocity υ_*f*_ in the 3D vascular network is calculated by solving the continuity and momentum equations in COMSOL Multiphysics. The flow is assumed to be laminar, incompressible, Newtonian and without magnetization. The inlets and outlets of vascular networks are predefined. Constant flow speed *u*_*in*_ is applied to the inlets, and zero pressure boundary condition is applied to the outlets. For the vessel walls, the non-slip boundary condition is applied.

The efficiency of magnetic steering is quantified by the total magnetic energy consumed by the particle during time interval *t* as 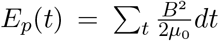 where *B* = *µ*_0_(*H* + *M*) is the magnetic flux density (magnitude of the magnetic field), where the magnetization of the fluid is neglected (i.e. *M* = 0).

### 2.3 Reconstruction of the vascular network

Artificial tree-like vascular networks and web-like vascular networks in 2D and 3D spaces are constructed. The length and diameter of vascular branches are selected according to Murray’s law [53], which minimizes the resistance to flow through the network when the network flow is smooth and leak free. Specifically, if a branch in vessel generation *j* which has a diameter *D*_*j*_ splits into *n*_*b*_ branches, the diameter of the *nb* branches in generation *j* +1 is *D* and can be calculated as 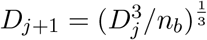

The length of the vessel branch *Lj* also follows Murray’s law 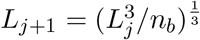 Based on Murray’s law, tree-like networks which have two splits and three splits are constructed. For each network, the length and diameter of the trunk (vessel branches in generation 1) are set as 1000 *µm* and 200 *µm* respectively. Each network contains four vessel generations. The values of *D*_*j*_ and *L*_*j*_ are shown in Table A1 in the Appendix. To construct an *in vivo* like vascular network, the network skeleton and diameters of vessel branches are extracted from a 2D fluorescent image of blood vessels [54].

Next, the 2D and 3D network model are built in Solidworks, and converted to triangle meshes in COMSOL Multiphysics. A laminar fluid flow study is performed in COMSOL. The mesh information are used to reconstruct the vascular structures and the 3D flow field using a “nearest” interpolation method in Python. The boundary layers are also defined accordingly.

### 2.4 Computational procedure

The numerical multi-physics model for particle dynamics in the vascular networks is solved using an Euler-Lagrange approach. First, the vascular structures, boundary layers and fluid flow field is constructed. High dimensional matrices that stores coordinates ***x*** = [*x, y, z*], boundary information ***M*** _*b*_, fluid velocity field ***M*** _*u*_, ***M*** _*v*_, ***M*** _*w*_, and the magnetic field information ***B*** and ∆***B*** are constructed. Next, the trajectory of a single magnetic particle in the network is calculated based on Algorithm 1. Specifically, to study the moving efficiency of RWSP, PRW and Lévy walk, the kinematic model discussed in section 2.1 is used. To develop an optimal magnetic steering strategy, the transient moving speed and direction of the particle are calculated based on the governing equations (i.e., Eqs. (3), (4), and (13)). The governing equations are solved using first-order Euler’s method in time, where the time step *dt* is set as 0.1 *s*.

The pseudo code of the multi-physics model is presented below.

#### Algorithm 1

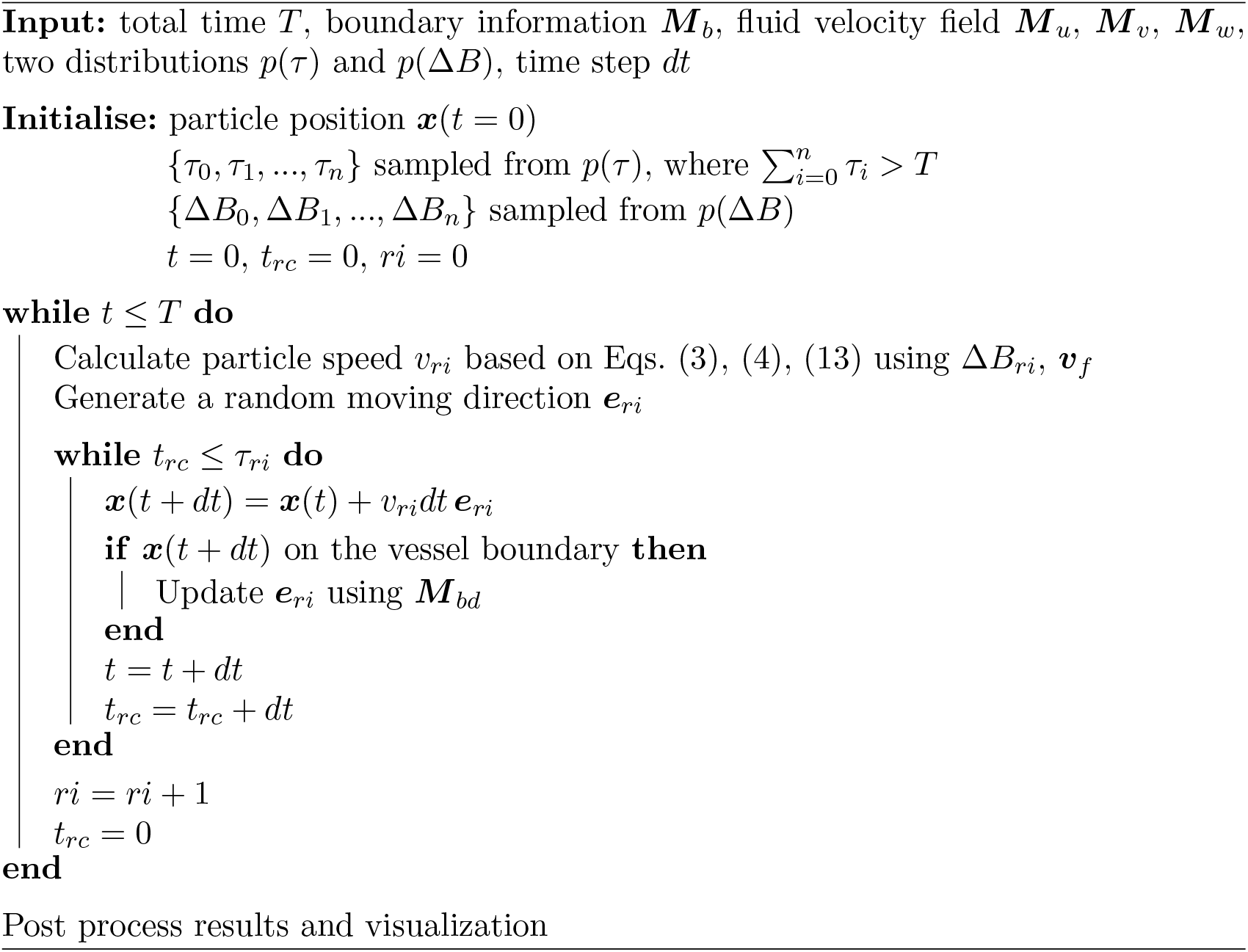

## 3. Results and discussion

### 3.1 Particle dynamics in 2D artificial networks

First of all, the dynamics of magnetic particles moving in 2D artificial vascular networks is analyzed. To investigate the effects of network topology on particle dynamics, we constructed three types of vascular networks, including bifurcation and trifurcation tree-like networks and a web-like network. The network structure is defined by its branch lengths (*L*_1_, *L*_2_, *L*_3_ and *L*_4_), diameters (*D*_1_, *D*_2_, *D*_3_ and *D*_4_), and the splitting angle *θ*, as shown in Figure 2 (a). Parameter values are listed in Table A1 in Appendix. The relationship between *L*_*j*_ and *L*_*j*+1_, *D*_*j*_ and *D*_*j*+1_ (*j ∈* [1, 2, 3]) follows Murray’s law (detailed discussions are provided in section 2.3). Figure 2 (b) shows four examples of the vascular networks, where particles move in the red areas. Examples of particle trajectories are shown in Figure 2 (c). Due to the large probability of having long directional persistence *τ*, particles performing RWSP and Lévy walk motion are likely to achieve long straight movements than PRW. RWSP has the largest mobility and particles are able to reach the ends of vascular branches in a short time period. Though many straight movements are truncated due to the reflection at vessel boundaries, the mean persistent length (i.e., length of straight movements) of RWSP is still larger than Lévy walk (Figure 2 (d)). Interestingly, we found that the topology of networks does not have a significant influence on particle mobility. For example, in the trifucation tree-like networks, the *α* exponent of MSDs (i.e., *MSD ∼ t*^*α*^) are similar when the splitting angle *θ* changes from 30^*◦*^ to 150^*◦*^ (Figure 2 (e)). But *α* is influenced by moving strategies. In particular, *α*_RWSP_ *> α*_Lévy_ *> α*_PRW_ *≈* 1. Figure 2 (f) shows the comparison of MSDs of Lévy walk and RWSP in a 60^*◦*^ bifurcation network, a 60^*◦*^ trifurcation network and a web-like network. The results suggest that particle mobility is less influenced by network structures, but RWSP is usually faster than Lévy walk in the vascular networks.

**Figure 2.**
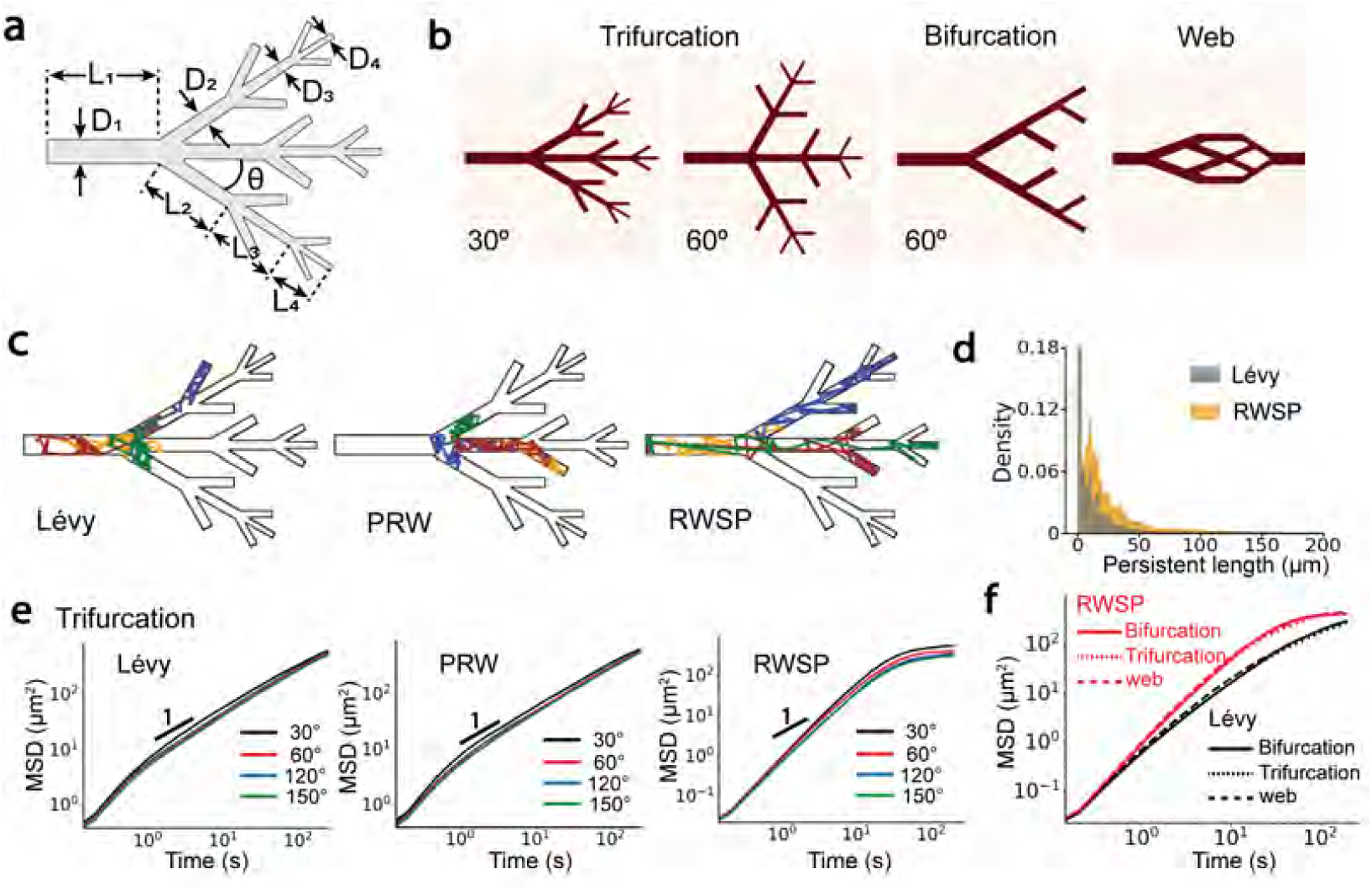
Particle dynamics in 2D artificial vascular networks. **(a)** Schematics of the 2D tree-like vascular network which has a trifurcation angle *θ*, branch lengths *L*_*j*_ and vessel diameters *D*_*j*_ (*j ∈* [1, 2, 3, 4]). **(b)** Examples of the trifurcation (*θ* = 30^*◦*^, 60^*◦*^) and bifurcation (*θ* = 60^*◦*^) tree-like networks and a web-like network. Areas inside the vessels are labeled as red. **(c)** Particle trajectories in the 60^*◦*^ trifurcation network where particles perform Lévy walk, PRW and RWSP motions. Different trajectories are labeled by different colors. **(d)** Distributions of persistent length when particles perform Lévy walk and RWSP motions. **(e)** MSDs of particles moving in the trifurcation tree-like networks influenced by the splitting angle *θ* and moving strategies (i.e., Lévy walk, PRW and RWSP). **(f)** MSDs of particles influenced by the vascular topology (i.e., bifurcation, trifurcation, web-like) and moving strategies (i.e., Lévy walk, RWSP).

### 3.2 Particle dynamics in 3D artificial networks

Next, particle dynamics in 3D artificial tree-like and web-like networks are investigated. Figure 3 (a) shows the example particle trajectories in a 60^*◦*^ trifurcation network. Like the 2D situations, particles that have RWSP type of motion are likely to take long straight movements and reach the far ends of the network in a short time, while particles diffuse locally in the PRW motion. Vessel boundaries have a larger effect on the Lévy walk motion in the 3D space, such that long movements become rare. By quantifying the persistent lengths of Lévy walk and RWSP, Figure 3 (b) shows that the average persistent length of RWSP is much larger than Lévy walk. Many long straight movements, whose lengths are larger than 30 *µm*, are maintained in RWSP. On the whole, persistent lengths in the 3D networks are larger than 2D networks regardless of the motion types. Moreover, the MSDs of particles are rarely influenced by the network topology. The MSD curves collapse onto the same curve when the splitting angle *θ* increases from 30^*◦*^ to 150^*◦*^ (Figure 3 (c)). The *α* exponent of MSDs are close to 1 for both Lévy walk and PRW, which indicates that the motion becomes diffusive. For RWSP, *α* is slight larger than 1. Figure 3 (d) further demonstrates that the dynamics is less influenced by the network structures as the MSDs are similar in bifurcation, trifurcation tree-like and web-like networks. But RWSP can also achieve faster mobility than Lévy walk in various 3D vessel netowrks.

**Figure 3.**
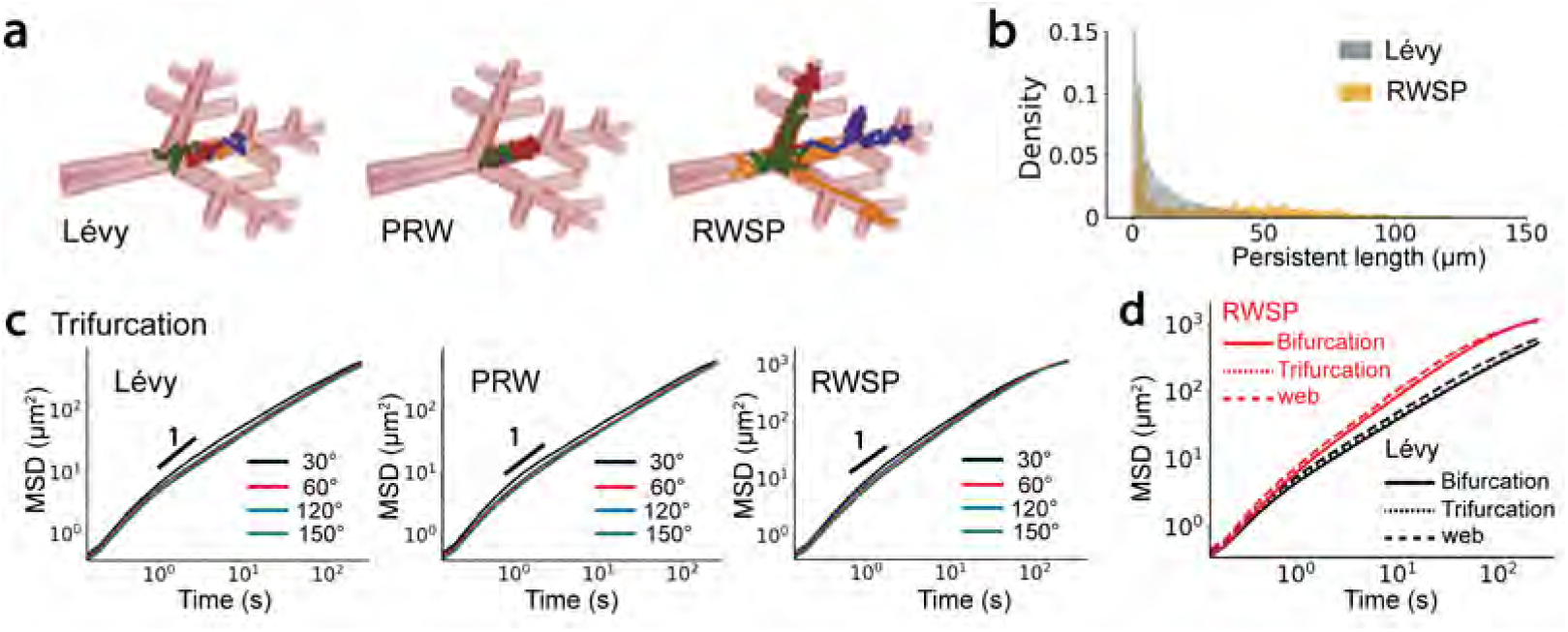
Particle dynamics in 3D artificial vascular networks. **(a)** Particle trajectories in the 3D trifurcation network (*θ* = 60^*◦*^) where particles perform Lévy walk, PRW and RWSP motions. Different trajectories are labeled by different colors. Vessel walls are labeled as red. **(d)** Distributions of persistent lengths when particles perform Lévy walk and RWSP motions. **(e)** MSDs of particles moving in the trifurcation tree-like networks influenced by the splitting angle *θ* (*θ ∈* [30^*◦*^, 60^*◦*^, 120^*◦*^, 150^*◦*^]) and moving strategies (i.e., Lévy walk, PRW and RWSP). **(f)** MSDs of particles influenced by the vascular topology (i.e., bifurcation, trifurcation, web-like) and the moving strategies (i.e., Lévy walk, RWSP).

### 3.3 Magnetic particle steering strategy

Based on the kinematic analysis of particle dynamics in the 2D and 3D vascular networks, results have showed that the RWSP type of motion can substantially improve the particle mobility. In order to manipulate particles to achieve the RWSP type motion, a time-varying magnetic field is used. In this section, the design principles of the stochastic magnetic steering strategy is firstly introduced. Then the particle motion controlled by the steering strategy in the *in vivo* like vascular network is simulated using the multi-physics model. Finally, the efficiency of several control strategies of the magnetic field is compared, and the optimal magnetic steering strategy is selected.

Firstly, to efficiently navigate in a complex vascular network without prior knowledge of the network topology and the current particle location, a time-varying stochastic steering strategy is used. Specifically, at the beginning, two series of random numbers *{τ }* = *{τ*_0_, *τ*_1_, …, *τ*_*n*_*}, {*∆*B}* = *{*∆*B*_0_, ∆*B*_1_, …, ∆*B*_*n*_*}* are sampled from two distributions *p*(*τ*) and *p*(∆*B*). In this study, 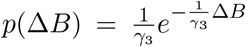 where *γ*_3_ is set as 10 *mT/m*. Four types of *p*(*τ*) are studied. Each corresponds to a steering strategy.

Specifically,

1. Constant: *τ* is a constant;
2. PRW: 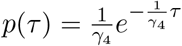
3. Lévy: *p*(*τ*) *∼ τ* ^*−β*^;
4. RWSP: 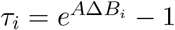 for *i ∈* [0,n]

At time points 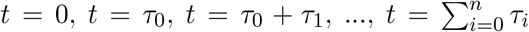 the magnetic field is updated. The direction of the magnetic field is randomly selected. The magnetic field at position ***x*** is calculated as *B*(***x***) = *B*_*c*_ + ∆*B*_*i*_||***x***||*/x*_*c*_, where ∆*B*_*i*_ *∈ {*∆*B}*. ||***x***|| is the norm of ***x***, and *x*_*c*_ is used for non-dimensionlization. *B*_*c*_ is a constant. In this study, *B*_*c*_ is set as 0.5 *T*. And the direction of the magnetic field is randomly selected. Note that, in order to satisfy the maximum limit of magnetic exposure of human body, less than 3 *T* permanent magnetic field is usually applied [27]. And in the state-of-art MRI techniques, the magnitude of the static magnetic field that can be generated is as high as 7 *T*. For the gradient magnetic field, 300 *mT/m* gradient can be achieved. Thus, the parameters used in this study are both achievable and reasonable [55].

Secondly, a multi-physics model is constructed to analyze the particle dynamics influenced by the vascular structure, magnetic field and fluid flow field. The workflow is shown in Figure 4. Specifically, the vessel structure is extracted from a fluorescent image of the deep vascular plexus of the retina [54]. The 3D vascular boundaries are reconstructed based on the segmentation information. Next, the fluid flow field inside the 3D vascular network is numerically solved. Then, the time-varying magnetic field and the gradient of the magnetic field are generated based on the steering strategies. Note that, only the open-loop control of the magnetic field is investigated here. Thus, the information of the magnetic field during the whole simulation time can be acquired before calculating the dynamical equations of the particle. The information of vascular boundaries, fluid flow field and magnetic field are stored as high-dimensional matrices. Finally, Eqs. (3), (4), (5), (13) are numerically solved using Euler’s method. Particle trajectories are visualized, and the MSDs (or the first passage time, FPT) are calculated and used to quantify the particle mobility. Furthermore, a grid search method is used to optimize the magnetic steering parameters based on the criteria of maximizing the exponent *α* of MSDs.

**Figure 4.**
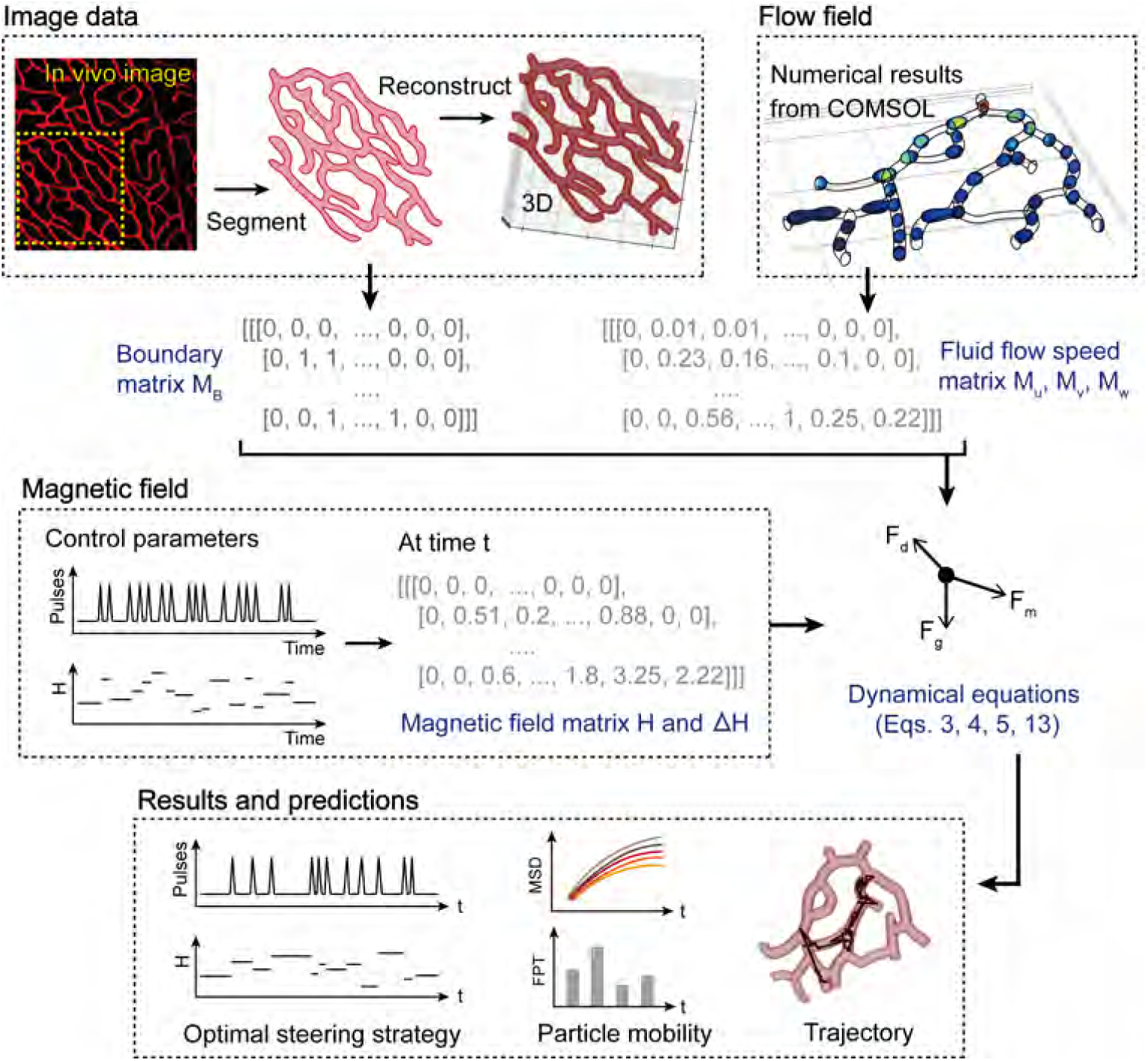
Schematics of the multi-physics model and the methodology used to study particle dynamics influenced by the 3D vascular boundaries, gradient magnetic field and fluid flow field.

An example of steering a magnetic particle using the stochastic magnetic field is shown in Figure 5. Pulse signals generated at time points 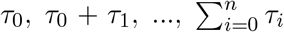 are shown in Figure 5 (a). The magnetic force experienced by the particle is shown in Figure 5 (b). The movements of the particle in the 3D vascular network at five different time points are plotted in Figure 5 (c).

**Figure 5.**
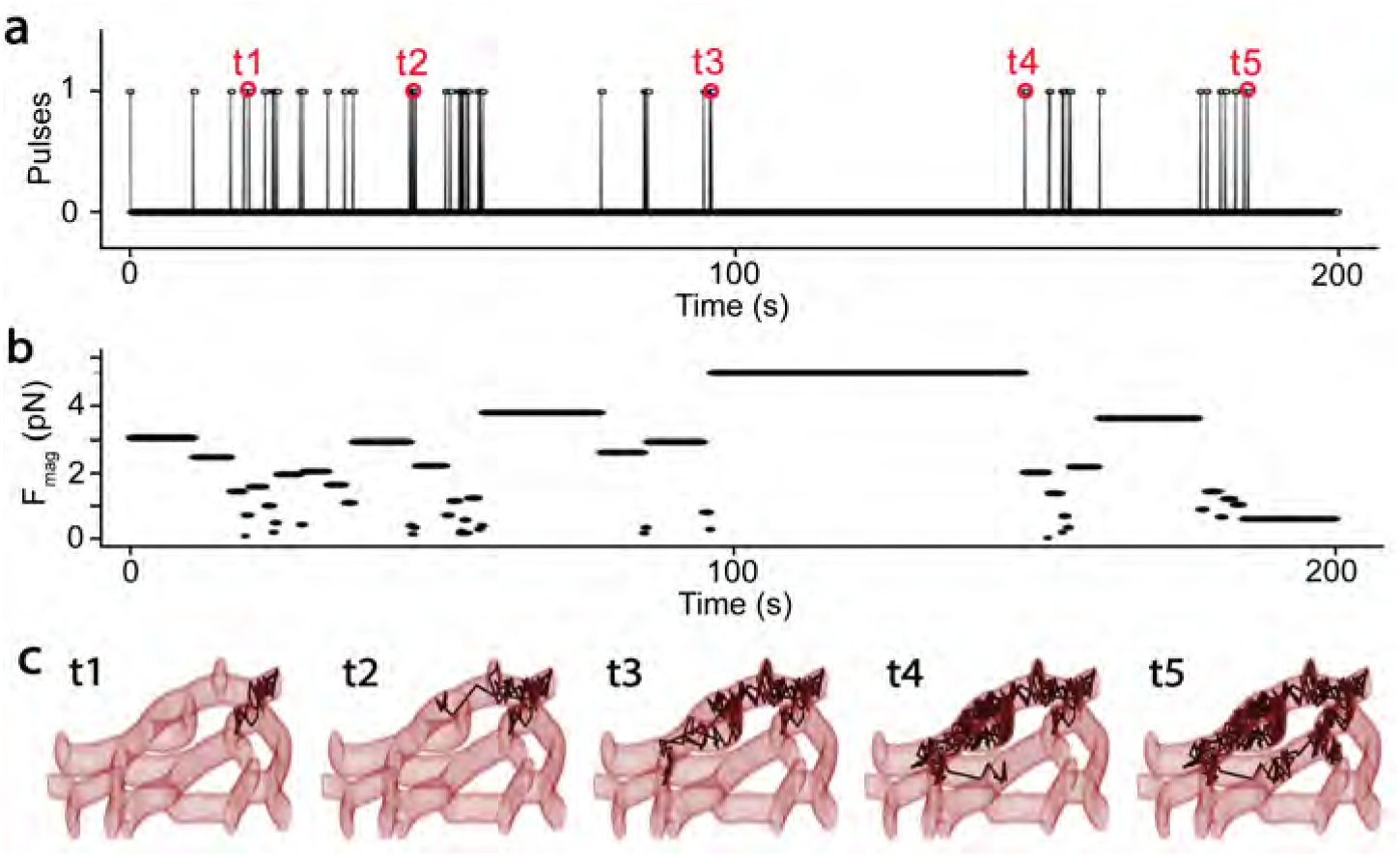
An example of magnetic steering of a particle in an *in vivo* like vascular network. **(a)** Pulse signals generate over time. When the value changes from 0 to 1, the direction and magnitude of the magnetic field are adjusted. **(b)** The magnitude of the magnetic force over time. **(c)** Snapshots of the movements of a magnetic particle in the vascular network. The snapshots are taken at five time points highlighted by red circles in **(a)**.

### 3.4 Efficiency of magnetic steering strategies

The efficiency of the four magnetic steering strategies introduced in section 3.3 and the influence of the steering parameters are examined here. The first strategy (“Constant”) changes the direction of magnetic field for every *τ* seconds. *τ* is a constant steering parameter which changes from 0.5 *s* to 20 *s*. The second “PRW” method changes the magnetic field after a time interval *τ*_*i*_, where *τ*_*i*_ follows an exponential distribution with a mean value *γ*_4_ changes from 0.5 *s* to 20 *s*. For the third “Lévy” strategy, the steering parameter *β* which determines the long-tailed distribution of *τ*_*i*_ changes from 1.1 to 2.9. And the steering parameter *A* in the fourth “RWSP” method changes from 0.5 to 20. ∆*B*_*i*_ follows an exponential distribution with a mean value *γ*_3_ equal to 10 *mT/m*. Note that, the influence of ∆*B* is not investigated here, because a large magnetic field applied a large force on the particles and will always increase the particle velocity and the MSDs. But in reality, there is a maximum limit of the magnetic field intensity that can be applied to the human body.

Examples of pulse signals generated to trigger the alteration of the magnetic field for the four steering strategies are shown in Figure 6 (a1), (b1), (c1) and (d1). As expected, the time interval between two pulses and the magnitude of the magnetic field are kept as constant for the “Constant” method. Magnetic field changes frequently and the time interval between two changes is relatively similar in the “PRW” method. Frequent changes and long-term persistence co-exist in the “Lévy” and “RWSP” methods. In the “PRW” method, the fast increasing of MSD is achieved when *γ*_4_ = 7.5 *s*. In the “Lévy” method, when *β* = 1.4, the largest MSD is achieved. In the “RWSP” method, the fast increasing of the MSD curve is achieved when the steering parameter *A* is around 10. And in the “Constant” method, *τ* = 10 *s* corresponds to the fast increasing of the MSD.

**Figure 6.**
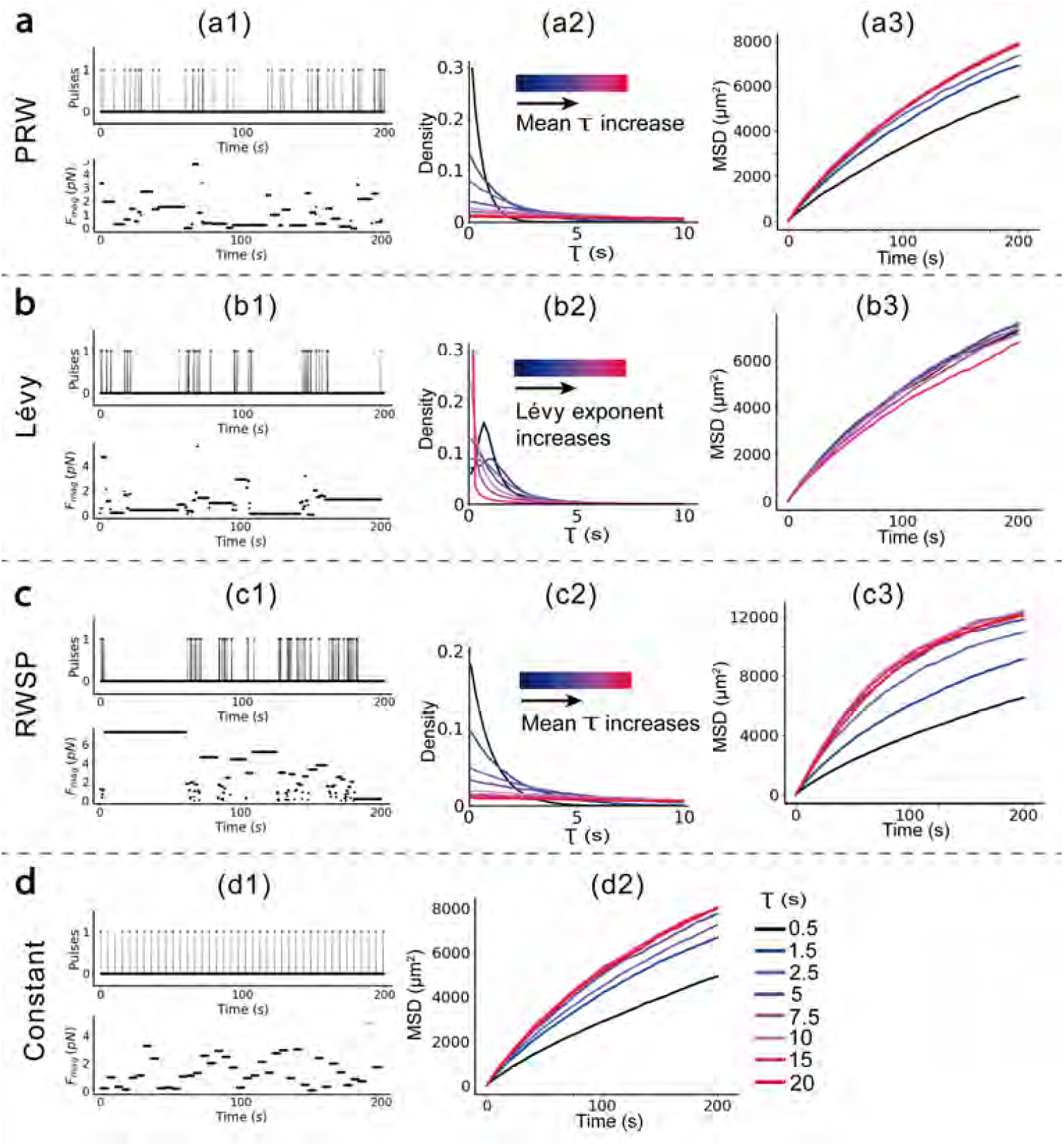
Parameter optimization and comparison of the four magnetic steering strategies. (a) A “PRW” steering strategy. (a1) The pulse signals and magnitudes of the magnetic force over time. (a2) Distributions of persistence *{τ }*. The mean persistence *γ*_4_ increase as the color changes from blue to red. (a3) MSDs influenced by the steering parameter *γ*_4_. **(b)** A “Lévy” steering strategy. (b1) The pulse signals and magnitudes of the magnetic force. (b2) Distributions of persistence *{τ }* influenced by Lévy exponent *β*. (b3) MSDs influenced by steering parameter *β*. **(c)** A “RWSP” steering strategy. (c1) The pulse signals and magnitudes of the magnetic force. (c2) Distributions of persistence *{τ }* influenced by the steering parameter *A*. The mean value of persistence increases as *A* increasing. (c3) MSDs influenced by the steering parameter *A*. Note that, for figures in the same row (e.g. (a2) and (a3)), the distribution and MSD curve have the same steering parameter if they are labeled with the same color. **(d)** A “constant” steering strategy. (d1) The pulse signals and magnitudes of the magnetic force on a particle. (d2) MSDs influenced by steering parameter *τ*.

Furthermore, the best parameters of the four steering strategies are identified based on a grid search optimization method. And the MSDs of particles and the magnetic energy consumed by the particles in 200 *s* of the four optimized magnetic steering strategies are compared. The results in Figure 7 shows that the “RWSP” strategy achieves the best performance. The exponent *α* of the MSD curve is larger than 1, which suggests the particle motion is superdiffusive. Due to the reflection of vessel boundaries, the particle mobility of other three methods are close to the diffusive motion at the long time scale. The energy efficiency of “RWSP” strategy is slightly worse than Lévy walk, but is better than “PRW” and “Constant” strategies.

**Figure 7.**
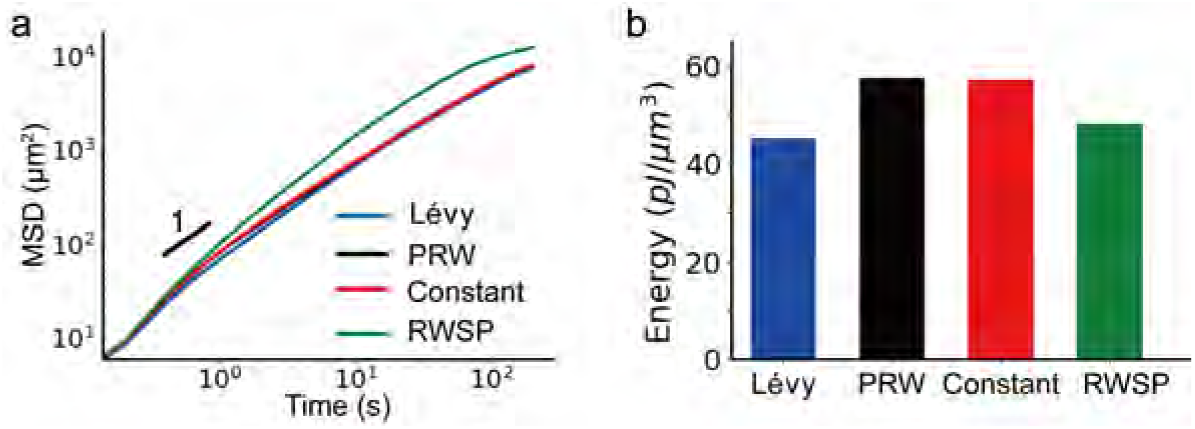
Comparison of the four types of steering strategies with optimized parameters. **(a)** MSDs of the particles manipulated by the four steering strategies with optimized parameters. **(b)** The total magnetic energy density consumed during the movement of the particle.

### 3.5 Particle steering influenced by heterogeneous flows

In the last part of this study, the influence of fluid flows inside the 3D vascular network on the particle dynamics is investigated. As shown in Figure 8 (a), fluid flows into the network through the two inlets and leaves the network from the five outlets. At the inlets, a constant fluid flow speed *u*_*in*_ is applied (*u*_*in*_ *∈* [5, 1000] *µm/s* is studied). Figure 8 (a) shows the fluid flow speed *u, v, w* in *x, y, z* directions when *u*_*in*_ = 50 *µm/s*. Due to the influence of the non-slip vessel walls, the flow speed along the *x* and *y* directions are highly heterogeneous. The flow speed in the three center branches which connect the inlets and the outlets 2, 3 and 5 is the largest. *u* and *v* can become as large as 100 *µm/s* in some places inside the vascular network. Particles are initially loaded at position P, which is shown by the black dot in Figure 8 (a). Particles are transported in the field depending on their position in the flow section. Particles on the edge move slower, and those in the center move faster due to the speed profile of the flow (Figure A1 (b) and (d)). When particles encounter the bifurcation, such as positions B1 and B2 shown in Figure 8 (a), depending on the flow conditions and geometry in particular, there is a division of the number of particles leaving on one branch and the other.

**Figure 8.**
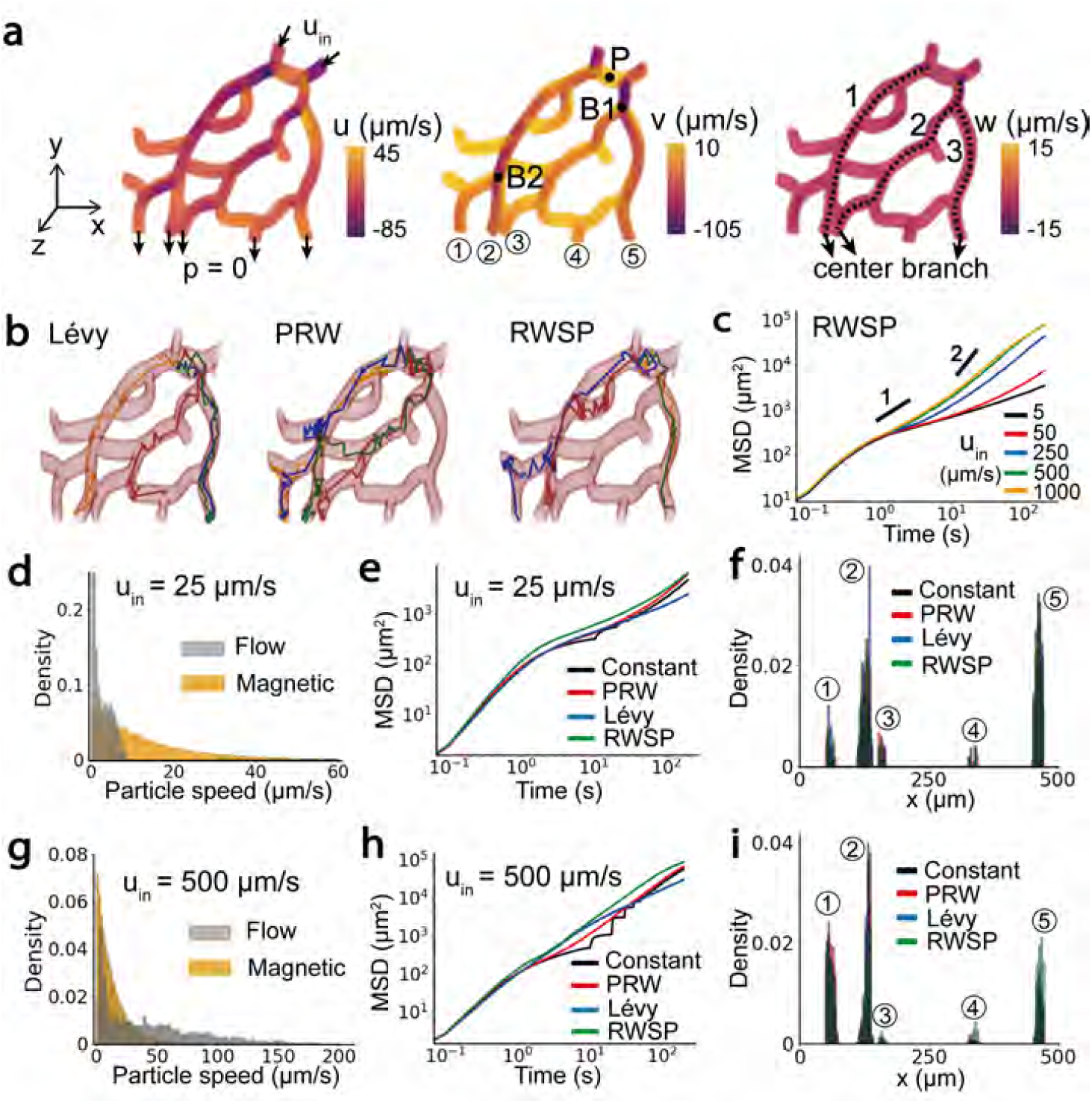
The dynamics of particles steered by the magnetic field influenced by heterogeneous flows inside the 3D vascular network. **(a)** The velocity components (*u, v, w*) of the flow field in the *x, y* and *z* directions. Negative values represent the fluid flows in the opposite directions. **(b)** Particle trajectories steered by the “PRW”, “Lévy” and “RWSP” strategies. For each type of motion, four trajectories starting from the same location P are represented by different colors. **(c)** MSDs of the particle steered by the “RWSP” strategy influenced by different inlet speed *u*_*in*_. At a small fluid flow speed when *u*_*in*_ = 25 *µm/s*, **(d)** distributions of particle speed generated by the hydrodynamic drag force and the magnetic force; **(e)** MSDs influenced by the four steering strategies; **(f)** distributions of particles at the five outlets. At a large fluid flow speed when *u*_*in*_ = 500 *µm/s*, **(g)** distributions of particles speed; **(h)** MSDs influenced by the steering strategies; **(i)** particle distributions at the outlets.

The dynamics of magnetic particles is influenced by the flow field. Specifically, particles tend to flow through the network from the three center branches which are shown by the dotted lines in Figure 8 (a). In the horizontal connecting branches, the flow speed is relatively small, and the effect of magnetic steering becomes considerable. Figure 8 (b) shows example trajectories of particles, where most trajectories align with the three central branches. Increasing the fluid flow speed inside the network by increasing *u*_*in*_, the slope of the MSDs at long times becomes larger (Figure 8 (c)).

When the flow speed is small (i.e., *u*_*in*_ *≤* 50 *µm/s*), the particle motion is subdiffusive due to the reflection at the vascular boundaries (the slope of the MSDs is smaller than 1). When the flow speed is large(i.e., *u*_*in*_ *≥* 250 *µm/s*), the particle motion becomes superdiffusive.

A detailed investigation is provided in Figure 8 (d)-(f) when *u*_*in*_ = 25 *µm/s* and (g)-(i) when *u*_*in*_ = 500 *µm/s*. Specifically, when *u*_*in*_ = 25 *µm/s*, the average moving speed of the particle generated by the hydrodynamic drag force is smaller than the speed generated by the magnetic force (Figure 8 (d)). When the “RWSP” steering strategy is applied, the MSD of the particle is slightly larger than the other three methods (Figure 8 (e)). These particles are more likely to be transported in the center branches 1 and 3, and accumulate at outlets 2 and 5 (Figure 8 (f), Figure A1 (a) and (b)). When *u*_*in*_ = 500 *µm/s*, the particle speed generated by the fluid drag force becomes larger than the magnetic force (Figure 8 (g)). The “RWSP” strategy still achieves the fast particle mobility (fast increase of MSD)(Figure 8 (h)). This time, more particles flow through the center branches 1 and 2, and accumulate at outlets 1 and 2 (Figure 8 (i), Figure A1 (c) and (d)). In addition, when the “RWSP” steering strategy is applied, particles at the three outlets (outlets 1, 2 and 5) tend to be more uniformly distributed for both *u*_*in*_ = 25 *µm/s* and *u*_*in*_ = 500 *µm/s* conditions.

## 4. Conclusion

Efficient delivering and controlling the release of therapeutic agents at the localized target sites has become a primary focus of medical and pharmaceutical research. Over the past few decades, lots of efforts have been spent to design biocompatible magnetism materials and devices to improve the drug targeting efficiency. The ability of magnetic delivery systems to navigate in the complex *in vivo* conditions is expected to open new avenues for disease theranostics such as curing cancers and artery diseases. In order to provide a better understanding of the dynamics of drug particles and design effective magnetic delivery strategies, in this work, a multi-physics dynamical model which simulates the transport of magnetic particles in the 3D vascular networks controlled by the dynamic magnetic field is developed. The mobility of particles influenced by the vascular topology, magnetic steering strategies, and the heterogeneous flow field is investigated. We found that particle motion is less influenced by the vascular topology but rather the characteristics of the microscopic particle movements. When the transient moving speed of the particle is correlated with its directional persistence in a nonlinear way (i.e., RWSP motion), particles are able to achieve fast spreading in both 2D and 3D vascular networks. The RWSP motion of magnetic drug particles in the vascular network is realized by applying a stochastic time-varying magnetic field. Based on the multi-physics model, parameters of a dynamic magnetic steering strategy are optimized and the delivery efficiency of particles manipulated by various magnetic steering strategies are compared. The multi-physics dynamical model is particularly designed to capture the interactions between particles controlled by the magnetic field and the complex *in vivo*-relevant network structures. It can also consider the influence of slowly flowing fluid on the particle dynamics. Thus, it is expected to apply the numerical model and method to a wide range of medical applications, such as drug delivery in solid tumors, retina and respiratory tract. For example, Figures 5-8 show the particle dynamics in vessel networks of deep plexus of human retina. The results demonstrate the following advantages of a time-varying gradient magnetic field in enhancing the spread and delivery of drug particles.

(1) The time-varying gradient magnetic field is capable of steering particles in tortuous tracts and networks, while a static magnetic field may guide particles to enter the dead ends.

(2) The energy efficiency of the time-varying magnetic field could be higher by properly selecting the control parameters of the magnetic field.

(3) The time-varying magnetic steering strategy can be adapted to complex, heterogeneous fluid environments.

(4) Rich particle motions can be generated for specific applications using the dynamic magnetic field.

In the future studies, the numerical model proposed in this work can be easily adapted to study more complex and realistic particle dynamics in the 3D vascular networks. For example, nanoparticles with a size range between 50 *nm* and 150 *nm* are widely used in drug targeting. Compared with the 5 *µm* particles studied in this work, the magnetic forces acting on the nanoparticles are reduced significantly. Based on Eqs. (3)-(5) and Eq. (13), one can easily find the relationship between particle speed and its radius as 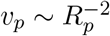 Thus, to steer nanoparticles in vascular networks, a large magnetic field is required. Carefully designing particle shape, material and material distribution (e.g., using multi-core magnetic particles) could potentially increase the magnitude of the magnetic force and improve the steering efficiency. The proposed numerical model can be extended to study the dynamics of magnetic particles of arbitrary shapes and components by modifying Eqs. (7)-(10) accordingly. Some previous studies have shown that small-size particles, especially nanoparticles, can be engulfed by the endothelial cells or penetrate through tissue layers [27, 50]. It would be interesting to carefully calibrate the particle dynamics near the vascular boundaries in the experiments. The leaky and slippery boundary conditions derived from the experimental measurements can be implemented into the model to improve the modeling accuracy. In addition, the influence of particle injection and releasing positions in the vascular network can to be further investigated using the model by adjusting the initial boundary conditions (the initial positions and speeds) of the particles. Moreover, previous studies have used 4-8 ordinary coils or coil arrays to generate static magnetic fields with many designable patterns and dynamic magnetic fields such as pulse and rotating fields [24]. Therefore, generating the gradient magnetic field in a random direction in discrete time points and realize the stochastic magnetic steering strategies discussed in this work is attainable.

To conclude, the multi-physics dynamical model and the results in this work can provide fundamental understanding and theoretical guidance in design and optimization of magnetic drug targeting systems. The novel stochastic magnetic steering strategy of particles in the 3D *in vivo* like vascular network requires only open-loop control of magnets, which is easy to implement in the experiments and can substantially improve the drug targeting efficiency. Thus, it is hoped to see a wide use of the stochastic steering strategy in the future drug delivery applications.

## Ethical Approval Statement

NA.

**Figure A1.**
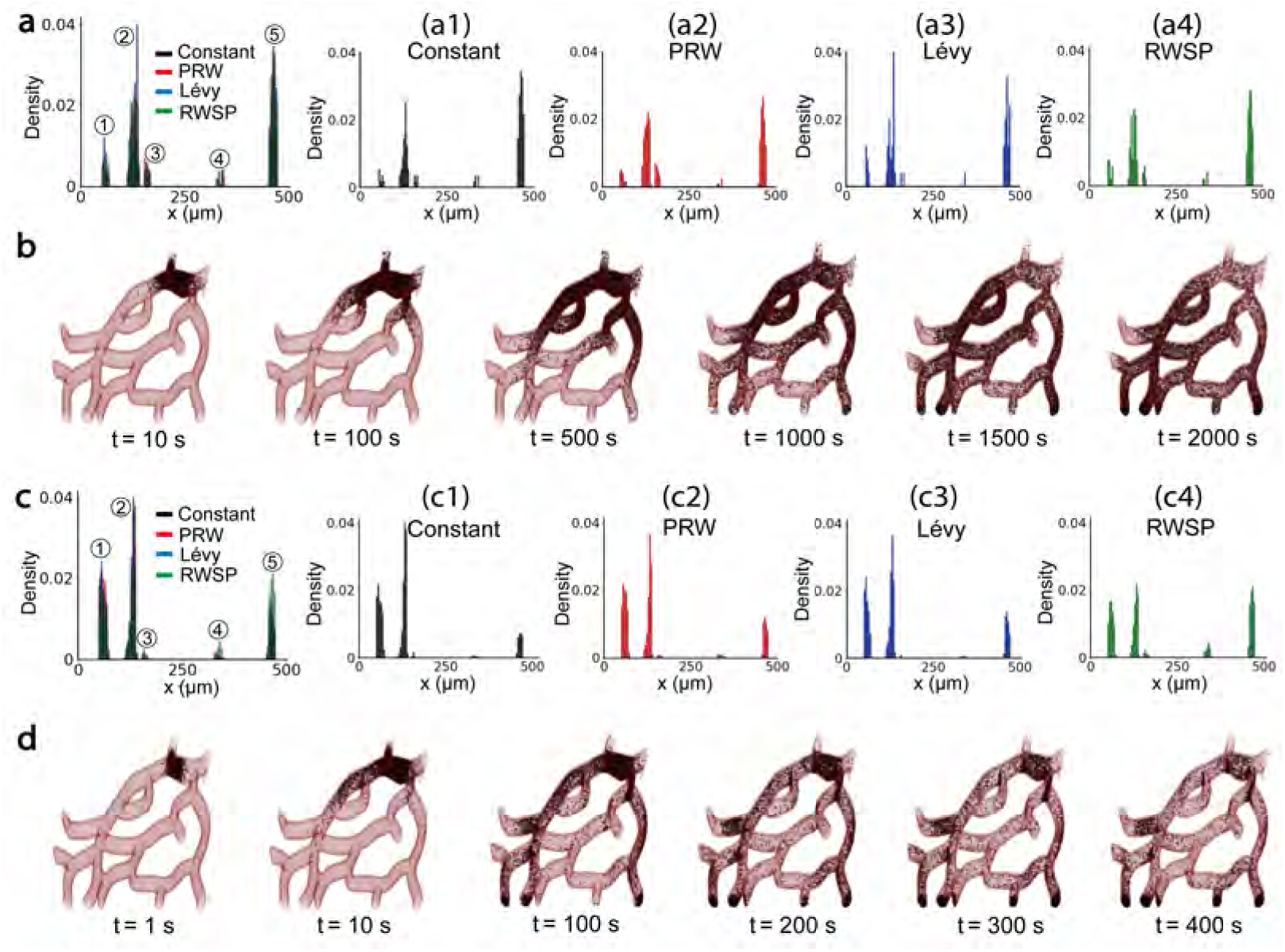
Distributions of particles. **(a)** When *u*_*in*_ = 25 *µm/s*, the distribution of particles at the five outlets influenced by steering strategies ((a1) Constant, (a2) PRW, (a3) Lévy, (a4) RWSP). **(b)** When *u*_*in*_ = 25 *µm/s*, particle distributions steered by the “RWSP” strategy at six time points. **(c)** When *u*_*in*_ = 500 *µm/s*, the distribution of particles at the five outlets influenced by steering strategies ((c1) Constant, (c2) PRW, (c3) Lévy, (c4) RWSP). **(d)** When *u*_*in*_ = 500 *µm/s*, particle distributions steered by the “RWSP” strategy at six time points.

## Data Availability

The code used to calculate particle motions steered by the magnetic field in the 3D vascular network is available at https://github.com/jessychen2018/Magnetic_Particle_Motion.

**Table A1.** Parameters of the 2D vascular network. Unit: *µm*

| | $L_1$ | $L_2$ | $L_3$ | $L_4$ | $D_1$ | $D_2$ | $D_3$ | $D_4$ |
| --- | --- | --- | --- | --- | --- | --- | --- | --- |
| 2 branches | 1000 | 794 | 630 | 500 | 200 | 159 | 126 | 100 |
| 3 branches | 1000 | 693 | 481 | 333 | 200 | 139 | 96 | 67 |

### Author Contributions Statement

Kejie Chen contributes to the conception, design, numerical analysis and drafting of the manuscript. Rongxin Zhou and Xiaorui Dong contribute to the numerical analysis, interpretation of the data and drafting of the manuscript.

